# Kinking and buckling instability in growing bacterial chains

**DOI:** 10.1101/2025.01.12.632655

**Authors:** Sean G. McMahon, John C. Neu, Jing Chen

## Abstract

Many gram-positive bacteria like *Bacillus subtilis* and *Clostridium* species, exhibit a growing chain–mediated sliding motility that is driven entirely by the force of cell growth. Particularly, the bacteria maintain cell-cell linkage after cell division and form long chains of many cells. The cells in a chain are continuously pushed outward by the mechanical force of cell growth. As the cell number in a chain grows, the cells toward the tip of the chain accelerate, and can in principle reach very high speeds. Although this seems to suggest a highly efficient motility mechanism, recent modeling work predicted mechanical stress builds up in the growing chain and the resulting chain breakage beyond a critical chain length, which ultimately sets a mechanical limitation in the maximum speed of the chain-mediated sliding. In this work we developed models to show that under different conditions the chain can either form sharp kinks or smooth buckles under the increasing stress. This can explain differential behaviors observed in different bacterial species. Our model further predicted how kinking and buckling affect the susceptibility of the chain to breakage. Our model provides a theoretical framework for predicting the dynamics and efficiency of growing chain–mediated bacterial sliding, and suggest cell properties that can optimize sliding efficiency.

## Introduction

Bacterial motility is necessary for bacterial cells to approach nutrients and avoid hazards, as well as for the bacterial colony to spread efficiently. Bacteria utilize a wide array of mechanisms to generate propulsion. Most of these mechanisms consist of dedicated motility appendages, such as flagella [1, 2] and type IV pili [3-5]. However, collective growth of sterically interacting bacterial cells can also generate expansive forces that push the cells forward and cause the colony to spread in space. This growth-driven motility is known as “sliding motility” [6-8]. Sliding motility has been well studied in the context of two-dimensional (2D) biofilm expansion, where *Bacillus subtilis* and *Vibrio cholerae* often serve as model organisms in both experimental and modeling research [9-16].

To gain efficiency over the basic 3D expansion of bacterial mass, bacterial sliding motility is often facilitated by additional mechanisms such as secreted surfactants or exopolysaccharides [6]. One of such facilitating mechanisms is the formation of long chains. In this type of sliding motility [4, 17-20], the bacteria maintain strong end-to-end connections between daughter cells following cell division; over multiple cell cycles, long and continuously growing cell chains form. Growth of individual cells forces the cells in a chain to move outward across the substrate. Through repeated cell division, the number of cells in the chain increases exponentially, potentially allowing the chain’s free tips to accelerate to very high speeds. Chain-mediated sliding motility has been observed in several bacterial groups, namely the *Clostridia* [4, 17-19] and *Bacilli* families [21-26], and is likely widespread among gram-positive bacteria as their thick cell wall can constitute a strong septum to maintain cell-cell connection after cell division.

Studying the mechanical dynamics of the growing chain is crucial for understanding the potential and biological implication of chain-mediated bacterial sliding. Our previous work, for example, predicted the mechanical limit of the chain-mediated sliding through mechanical modeling [20]. Particularly, our model shows that the mechanical stress builds up in the growing bacterial chain. Beyond a critical stress, the chain experiences mechanical instability, manifested as kinking that soon leads to chain breakage. The stress-induced breakage limits the maximum efficiency of bacterial colony spreading mediated by this sliding mechanism.

In this work we demonstrate through mechanical modeling that, in addition to the kinking instability, the growing bacterial chain can also experience mechanical instability as a smooth buckling of the chain. Our new continuum mechanics model predicts the critical condition for buckling instability. Comparing the result to the critical condition for kinking instability predicted by our previous work [20], we find that buckling is favored by low degree of anisotropy in cell-substrate drag, low overall cell-substrate drag, stiff cell-cell linkage, short cells, and/or lower cell growth rate. Furthermore, our chain simulations show that buckling reduces the mechanical stress in the chain and hence can delay chain breakage. However, buckling does not necessarily improve overall sliding efficiency, as the direction of chain growth is less persistent. Interestingly, the maximum sliding efficiency is found to happen at the transitional parameter region between buckling and kinking. Overall, our models shed light on the biological role of mechanical instability in the chain and potential strategies that a bacterium could exploit to optimize its chain-mediated sliding.

## Methods

### Continuum modeling of a growing bacterial chain

To predict conditions for buckling emergence, we develop a continuum mechanics framework for a growing chain with two free tips. In this continuum framework, we consider the following assumptions:

1. The chain grows uniformly at an exponential rate *r*. That is, the length of the chain grows in time as *L*(*t*) = *L*(0)*e*^*rt*^. This is to characterize continuous cell growth and exponential increase of cell number in the chain.
2. The chain has a uniform bending modulus *m*.
3. Each material point in the chain experiences a linear, anisotropic drag force from the substrate with a parallel drag coefficient *μ* and a perpendicular drag coefficient *ν*.

These assumptions capture the growth, bending, and viscous dynamics for the bacterial chains, especially in the limit where the persistent length of the chain is much larger than the typical cell length *l*_0_. The parameters of the continuum mechanics framework are summarized in Table 1.

**Table 1:** Physical parameters of the chain models.

| Continuum model | Rigid rod model |
| --- | --- |
| Typical cell length $l_0$ | Daughter cell length $l_0$ |
| Chain growth rate $r$ | Cell growth rate $r$ |
| Bending modulus $m$ | Angular spring constant $k_b$ |
| Parallel drag coefficient per unit length $\mu$ | |
| Perpendicular drag coefficient per unit length $\nu$ | |

Analysis of the continuum mechanics chain model yields a partial differential equation (PDE) for the curvature in the chain with respect to *ζ* and time *t* (Supplementary Materials). Non-dimensionalization of this PDE reveals that the dynamics of the chain depends only on two dimensionless parameters,

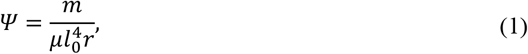

and

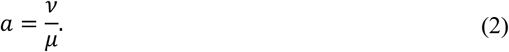

*Ψ* in Eq. (1) is the ratio between several physical parameters summarized in Table 1, and *a* in Eq. (2) is the anisotropic ratio between the perpendicular drag coefficient *ν* and parallel drag coefficient *μ. Ψ* and *a* are the only free parameters of the model. In the following sections, we analyze the PDE governing the curvature dynamics to quantify the emergence of buckling instabilities in the growing cell chain system and consider the biological implications of these dynamics.

### Modeling growing bacterial cell chains with discrete cells

For analyses that require a description with individual cells connected end-to-end, we use the discrete rigid-rod model introduced in [20] (Fig. 1B). In this model, individual cells are assumed to be inflexible rods with a time-dependent length, *l*(*t*) = *l*_0_*e*^*rt*^, which halves upon cell division. For simplicity of the simulation, cell divisions are assumed to occur synchronously when the cell doubles its length at birth *l*_0_. This simplification does not affect the overall mechanical dynamics of the cell chain. The cells are constrained by one another through physical connections. Bending of the chain is modeled by angular springs between adjacent cells with potential energy 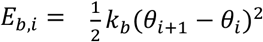. Like in the continuum framework, the cells experience an anisotropic translational drag force with parallel and perpendicular components. The cells also experience a rotational torque. Because the torque sources from the summed perpendicular drag along the length of a rotating cell, the rotational drag coefficient is coupled to the perpendicular translation drag coefficient by *ϵ* = *ν*/12 [20]. The rigid-rod model is formulated by a series of ordinary differential equations (ODE), which were used to predict the critical condition for kinking emergence [20]. In this work we will also use the rigid-rod model to simulate the chain dynamics. A more detailed description of the rigid-rod model, as well as derivations of the equations, can be found in the Supplemental Information of [20].

**Figure 1:**
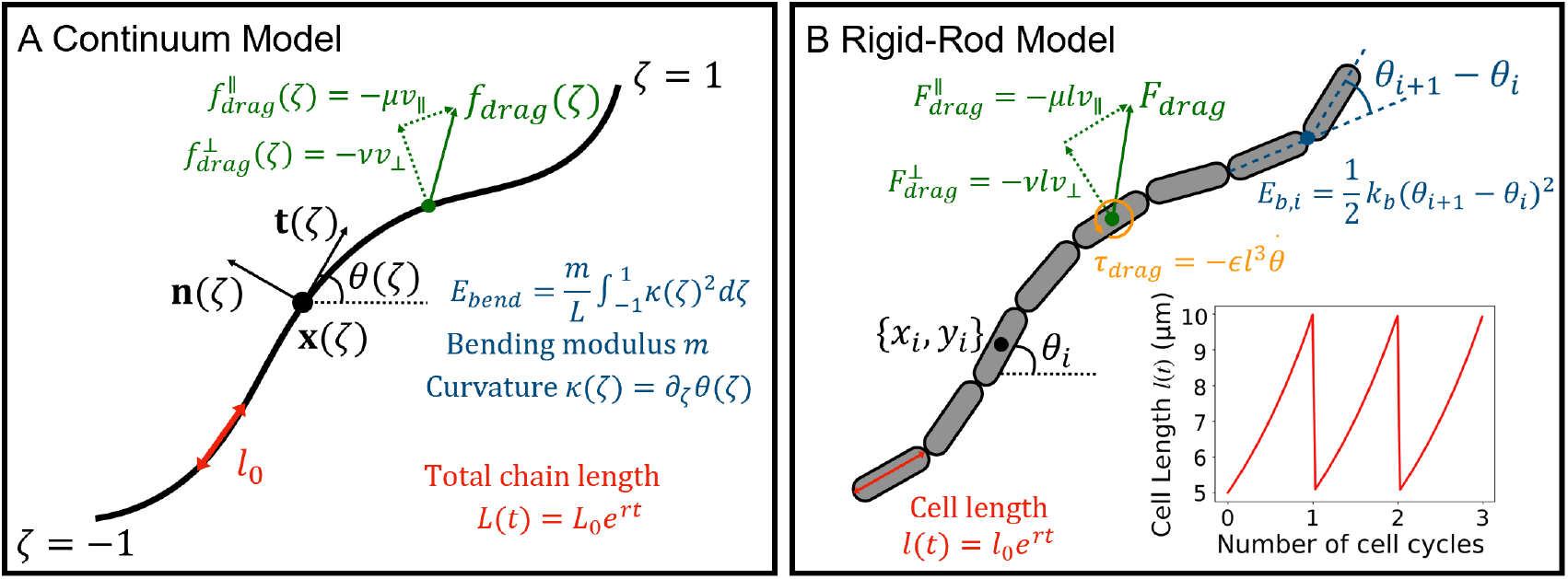
Mathematical models for growing bacterial cell chains. (A) Continuum mechanics model. *ζ* denotes the relative position of a material point along the arc length of the chain, such that −1 ≤ *ζ* ≤ 1 with *ζ* = −1,1 corresponding to the free tips of the chain. Each material point is associated with the coordinate relative to ground x(*ζ*), the unit tangent vector *t*(*ζ*) oriented in the direction of increasing *ζ*, and the unit normal vector n(*ζ*) defined counterclockwise with respect to the tangent. The angle between the tangent vector and the horizontal axis is given by *θ*(*ζ*). The chain grows exponentially with rate *rr* such that the total chain length is given by *L*(*t*) = *L*_0_*e*^*rt*^. The chain has a uniform bending modulus *m* and experiences a linear anisotropic drag from the substrate. (B) Discrete rigid-rod model. The chain is composed of discrete inflexible cells described by their center of mass coordinates {*x*_*i*_, *y*_*i*_} and angle from the horizontal axis *θ*_*i*_. The length of each cell grows in time as *l*(*t*) = *l*_0_*e*^*rt*^, and halves upon cell division (inset). Bending is implemented at the linkages between adjacent cells by angular springs with spring constant *k*_*b*_. The cells experience a linear anisotropic drag force and a linear rotational drag force from the substrate.

The physical parameters of the rigid-rod model correspond to those in the continuum mechanics framework one by one (Table 1). In the rigid-rod model, *l*_0_ is the cell length at birth. In the continuum framework the same *l*_0_ defines the system’s characteristic length scale. The rigid-rod model’s angular spring constant *k*_*b*_ is related to the continuous bending modulus *m* by *m* = *k*_*b*_*l*_0_. The parallel and perpendicular drag coefficients, *μ* and *ν* respectively, are the same between the two models. Therefore, like the continuum mechanics model, the discrete rigid-rod model also depends on only two free parameters, 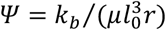 and *a* = *ν*/*μ. a* is identical between the two models, *Ψ* is equivalent after applying *m* = *k*_*b*_*l*_0_.

## Results

### Growing cell chains exhibit two modes of mechanical instabilities

Growing bacterial chains often exhibit one of two mechanical instability dynamics. The first instability is kinking which manifests as a sharp, localized bending in a cell-cell linkage as shown in the *C. perfringens* chain in Fig. 2A. Kinking has been observed in both the *Clostridia* and *Bacilli* families [20]. The kinked cell-cell linkage will soon snap, causing the chain to be broken in two. Mathematical models have examined the morphological changes in the chain driven by kinking emergence [25] as well the mechanical and population effects of cell linkage failure [20]. The second mode of instability is buckling, a response in the chain due to drag-induced stresses throughout the chain. Buckling has been well-studied in the context of Euler’s loaded beam. Conceptually, buckling in growing cell chains is similar to that in Euler’s beam: the large stresses cause the system to curve away from a straight configuration. However, the analogy is not perfect as bacterial cell chains are constantly growing and have discrete weak points at the linkages between adjacent cells where the chain is more susceptible to bending. The phenomenon of buckling in bacterial chains has been known for some time. Buckling, bending, and folding has repeatedly been observed in *B. subtilis* chains [24, 26, 27] and many mathematical modeling investigations have examined the morphological changes in these chains as a result of buckling [21-25, 28]. Buckling results in the emergence of a smooth curvature manifesting throughout large portions of the chain as in the *B. subtilis* chain shown in Fig. 2B.

**Figure 2:**
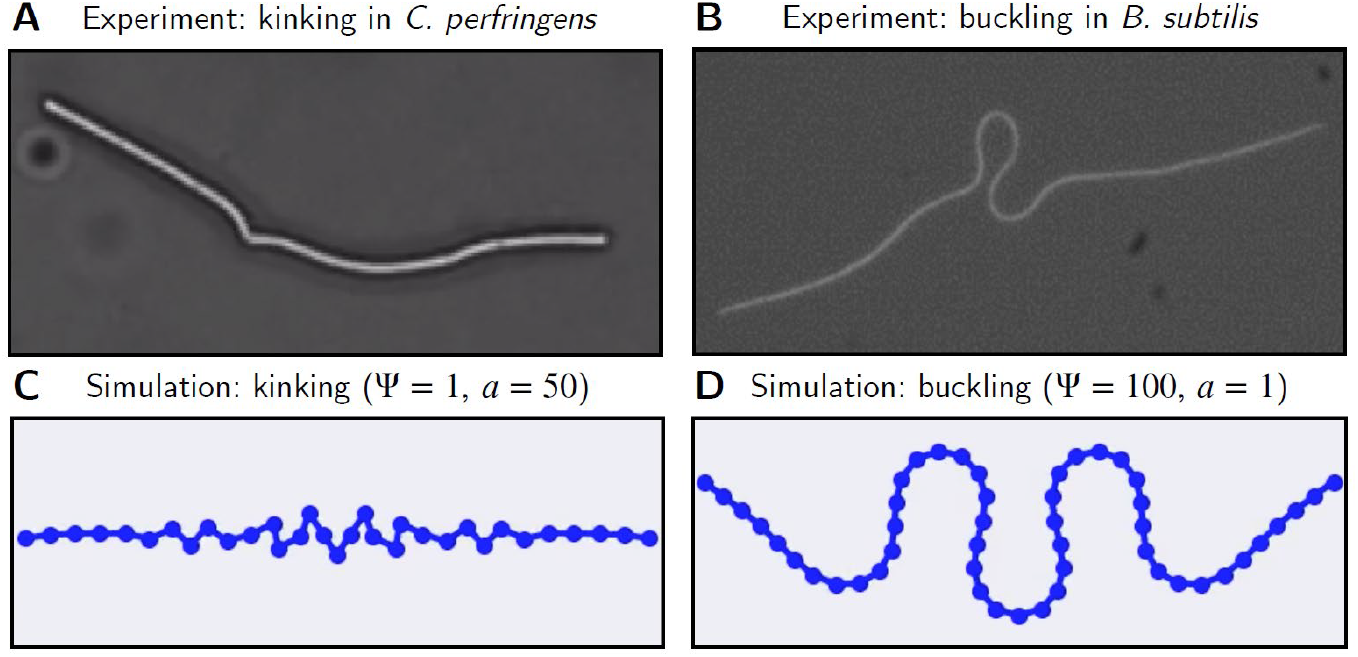
Kinking and buckling instability dynamics are observed both experimentally and in simulation. (A) A *C. perfingens* chain that grew on agar surface. After a period of mostly straight growth, a cell-cell linkage near the center of the chain began to kink. (B) A *B. subtilis* chain that buckled resulting in the emergence of large curvatures. Image cropped from Fig. 1b in [24] under a Creative Commons Attribution 4.0 International License. (C) A growing cell chain simulated using the rigid-rod model exhibiting the kinking instability for *Ψ* = 1, *a* = 50. (D) A growing cell chain simulated using the rigid-rod model exhibiting the buckling instability for *Ψ* = 100, *a* = 1.

Despite many observations of both the kinking and buckling instabilities, the physical conditions driving the emergence of one instability mode over the other, remain unclear. We simulated the chain dynamics using the rigid-rod model to examine which instability dynamic is dominant for different parameter values. These simulations confirm the emergence of both the kinking and buckling instabilities. Fig. 2C and D show simulation snapshots in which the chain exhibits typical examples of kinking and buckling, respectively. We see once again the kinking behavior is characterized by sharp bending in the cell-cell linkages. In Fig. 2C many linkages have bent sharply. In reality, a chain would likely break before multiple linkages bend sharply. The buckling observed in Fig. 2D is a smoother behavior that results in curvature emergence across multiple cells. The behaviors observed in the simulations in Fig. 2C and D qualitatively match the respective experimental observations shown in Fig. 2A and B.

Both the continuum mechanics and rigid rod models suggest the mechanical dynamics depend on only two parameters, *Ψ*, the dimensionless ratio between nearly all key physical parameters, and *a*, the anisotropic ratio of perpendicular to parallel drag coefficient. Since these two parameters dictate the model behaviors, we vary these parameters in order to observe the various dynamics that emerge across the *Ψ*–*a* phase space. Fig. 3 shows a series of snapshots from simulations with varied values of *Ψ* and *a*. Qualitatively, we observe that kinking is favored for small *Ψ* and large *a*, while buckling is favored for large *Ψ* and small *a*. One possible interpretation is that large values of *a* prevent dynamics with a longer length scale and hence kinking is favored. In the following sections, we will quantify the instability dynamic predictions across the *Ψ*–*a* phase space and consider the mechanical and biological implications of instability emergence.

**Figure 3:**
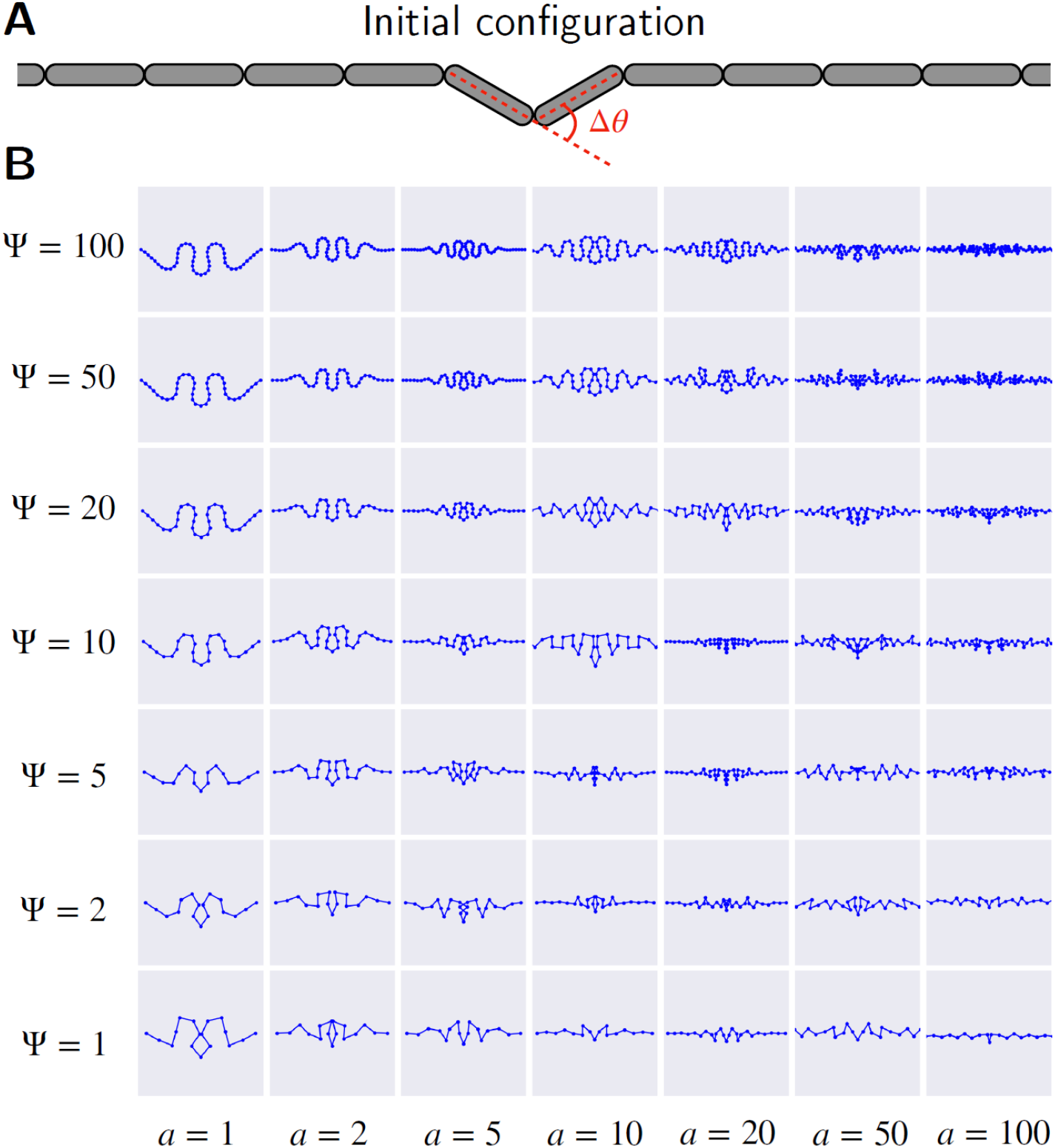
Mechanical instability dynamics in growing cell chains varies across the *Ψ, a* phase space. (A) The initial configuration used in all simulations. The chain is initialized with all cells perfectly aligned with the horizontal *x*-axis except for the two centermost cells, which are symmetrically perturbed away from the *x*-axis and make an angle of *Δθ* = 2^°^. (B) The configurations of bacterial chains simulated using the rigid-rod model for various values of *Ψ* and *a* after 100 minutes of simulation time. The dots correspond to linkages between adjacent cells. The chains have been initialized with various lengths chosen based on the expected dynamics. The simulated chains exhibit both sharp kinks and smooth buckling as well as behavior between the two. Some images have been cropped in order to more clearly highlight the instability dynamics in the center of the chain. The length scales vary across simulations. The full video of these simulations is included as Supplementary Movie 1.

### Perturbation analysis predicts critical chain length for emergence of buckling instability

In our previous work, we performed perturbation analysis on the discrete rigid-rod model (Fig. 1B) to derive the critical length of chain upon which kinking instability emerges and the chain becomes prone to breakage [20]. In that analysis, a small perturbation to the perfectly straight chain in the form of a small angle at the center cell-cell linkage. A chain above the critical length experiences high stress at the perturbed linkage, causing the angle of the linkage to increase and kinking to exacerbate. In contrast, in a chain below the critical length, the angle shrinks over time and the chain returns toward the perfectly straight shape. Therefore, the critical length is predicted by the point of sign change in the time derivative of the perturbed angle. Here we seek to derive the critical condition for buckling instability using a similar perturbation analysis. However, since buckling manifests as large regions of significant curvatures (Figs. 2D and 3), we can no longer focus on a single cell-cell linkage. A perturbation throughout the chain is needed and a continuum mechanics model (Fig. 1A) is best suited for this perturbation analysis.

In the perturbation analysis, we again consider small perturbations to a perfectly straight chain in the continuum model. This time the perturbation is assumed to be any arbitrary curving shape that is of small magnitude and satisfies the boundary conditions for the mechanical dynamics of the chain. Linearization of the weakly perturbed chain dynamics yields the nondimensionalized PDE for the time-dependent curvature *κ*(*ζ, t*) in the growing cell chain,

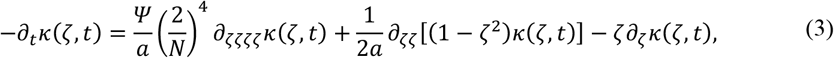

constrained by zero-force boundary conditions at the two free tips of the chain,

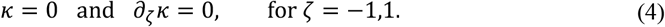

Buckling instability is defined as increase of the overall curvature of the chain over time after the initial perturbation. Here we quantify the overall curvature of the chain as

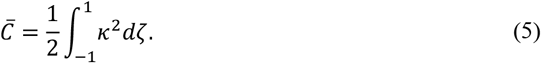

Combining Eqs. (3) and (5) yields

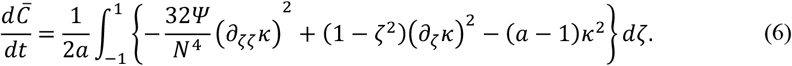

The full derivation is provided in the Supplementary Materials.

It is clear to see from Eq. (6) that increasing of the relative chain length *N* = *L*(*t*)/*l*_0_ causes increase in 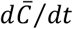. Therefore, there could exist a critical *N*_*crit*_ such that 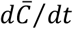 crosses zero, marking the point when buckling instability emerges. However, since the initial perturbation *κ*(*ζ*, 0) can be an arbitrary function, stability of the unperturbed, perfectly straight state, is only achieved when 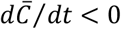 for all *κ*(*ζ*, 0). In other words, instability emerges when the integration in Eq. (6) stops being negative-definite. This cannot be solved in a closed form. We hence seek an approximate solution by decomposing the perturbation curvature profile *κ*(*ζ*) into a series of base buckling modes (Eq. (7)) that are similar to the chain shapes predicted by our rigid-rod simulation (Fig. 3). We can then plug these base buckling modes to Eq. (6) and determine the sign of 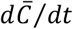 for each mode. Buckling instability emerges when any 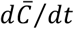 turns positive.

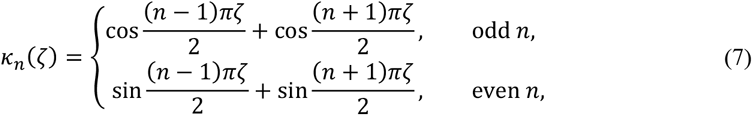

where *n* is the index for each buckling mode. Note that the buckling modes in Eq. (7) all satisfy the boundary conditions in Eq. (4). Fig. 4 shows the curvature profile for the first six buckling modes. Buckling modes with odd *n* are symmetric, while the mode with even *n* are asymmetric. All simulations consider in this investigation use the symmetric initial configuration shown in Fig. 3A and therefore our simulations only exhibit the symmetric buckling modes.

**Figure 4:**
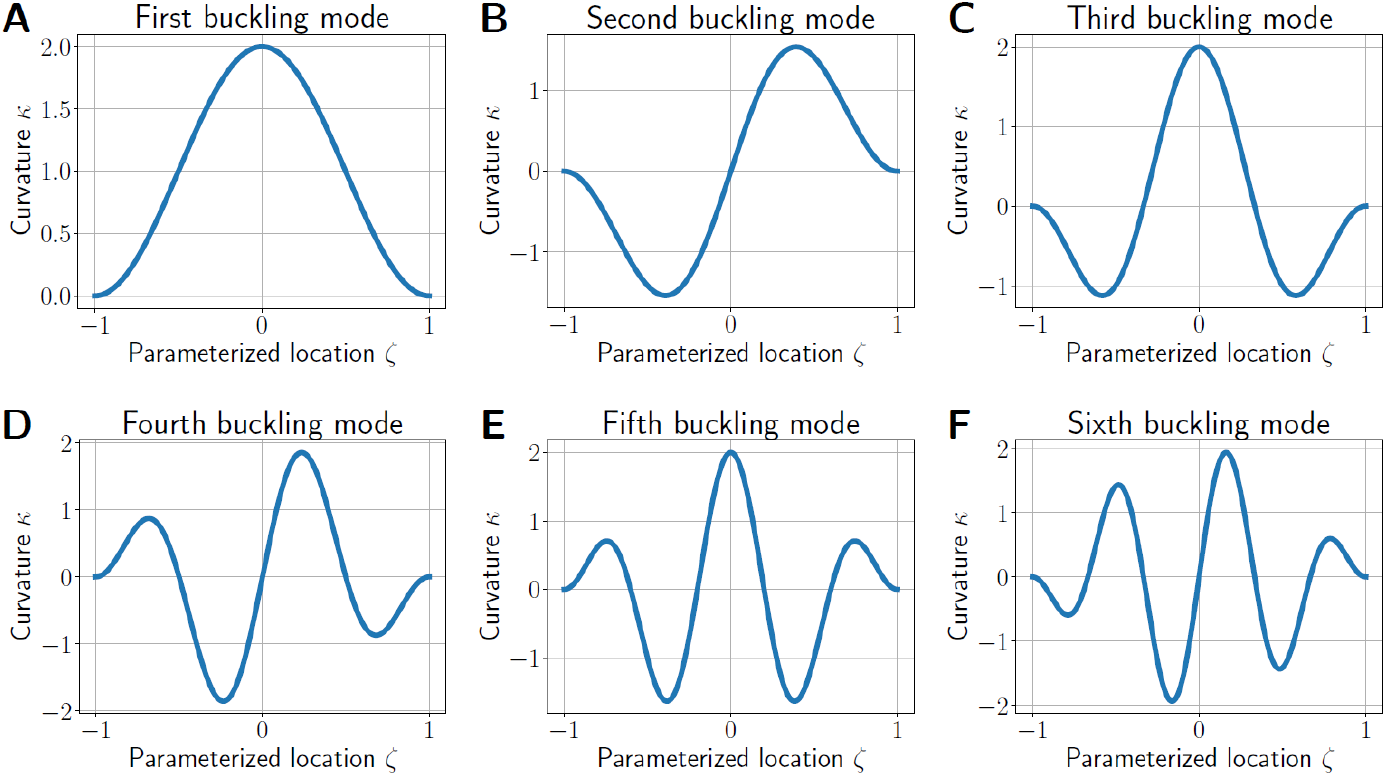
Curvature profiles of the first 6 buckling modes. The curvature profiles are plotted as a function of *ζ* according to Eq. (7). For the symmetric buckling modes (A), (C), and (E), the curvature is largest near the center of the chain then decreases in subsequent peaks before dropping to zero at *ζ* = −1,1 in accordance with the boundary conditions. In the asymmetric buckling modes (B), (D), and (F), the curvature is zero in the center of the chain and surrounded by peaks of decreasing magnitude before the curvature drops to zero at *ζ* = −1,1 in accordance with the boundary conditions. The derivative of the curvature is zero at the boundaries for all buckling modes provided by the ansatz. Note: for these plots, the *y* axes are not normalized to a meaningful physical scale. These are purely meant to represent the shapes of the curvature profile in the proposed buckling modes.

Through the calculation described above, the critical length associated with emergence of the *n*-th buckling mode is given by

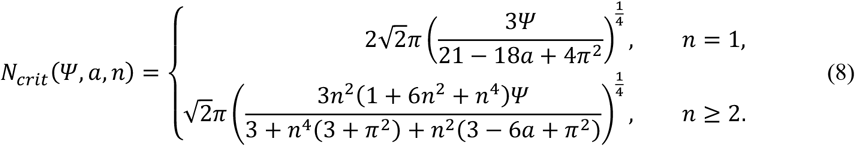

These expressions have been simplified under the assumption *n* is a positive integer. The expression for *n* > 1 is not valid for *n* = 1 because of a singularity in the expression prior to the application of the condition that *n* be an integer. Proper limit analysis results in the expression provided for *n* = 1.

Eq. (8) reveals a few interesting dynamics regarding buckling emergence. First, the relationship between *N*_*crit*_ and *n* is highly nonlinear which sometimes results in higher buckling modes emerging at shorter critical buckling lengths than lower modes (Fig. 5). Second, each buckling mode has a critical value of the anisotropic ratio *a* above which the critical buckling length becomes undefined, i.e., no buckling pattern in this mode expected. These critical *a* values are found by setting the denominators in Eq. (8) equal to zero and solving for *a*. The critical values of *a* as a function of the buckling mode *n* are

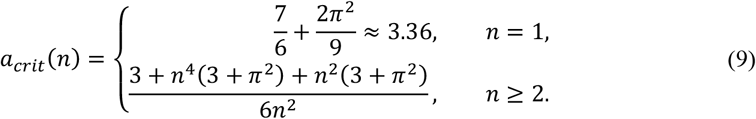

**Figure 5:**
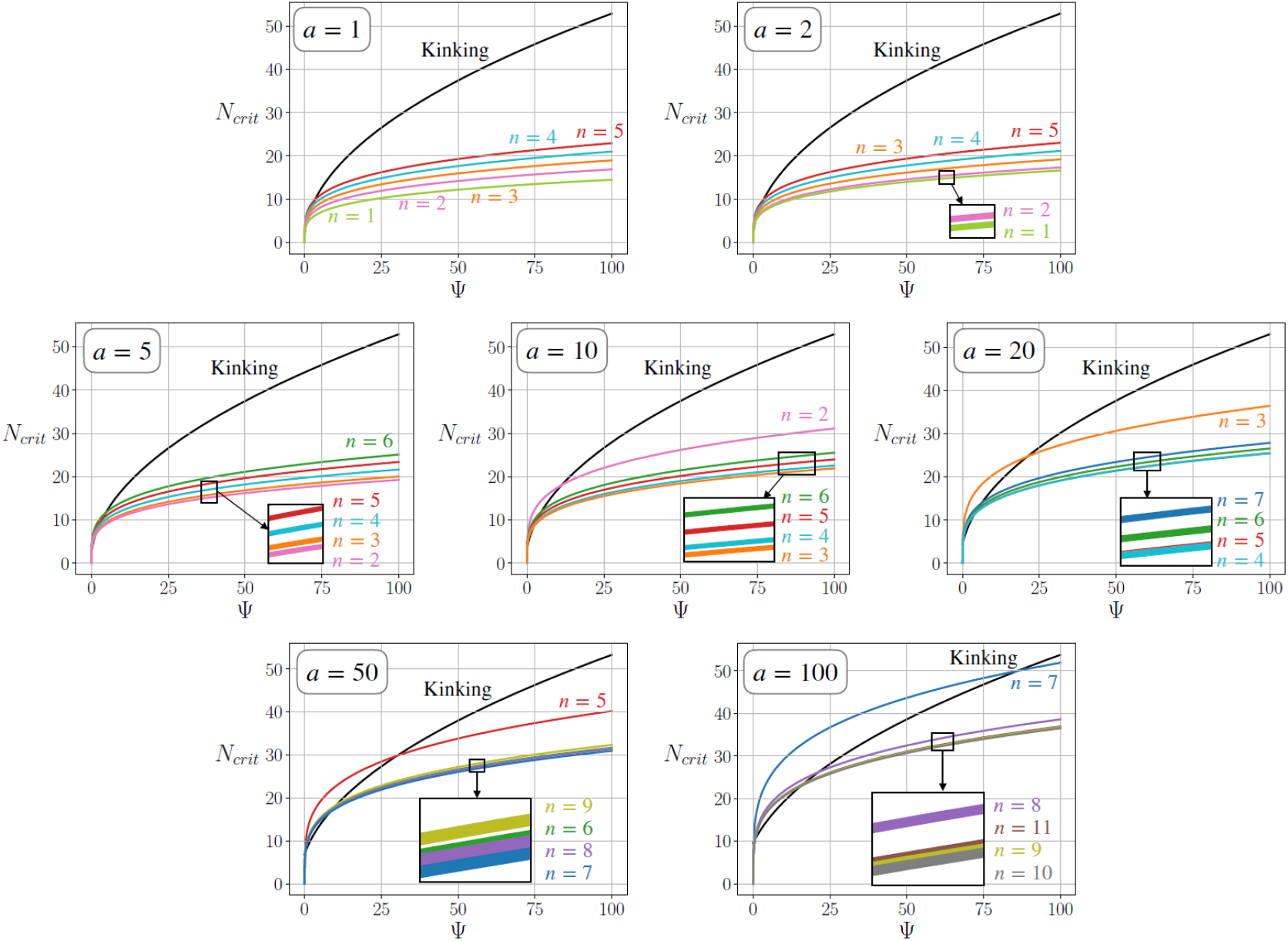
Predicted critical lengths for mechanical instability emergence. Plots of the critical buckling and kinking length as a function of *Ψ* for selected values of *a*. The critical buckling lengths are calculated using Eq. (8) and the critical kinking length are calculated from Eq. (10). Insets highlight the critical lengths when multiple modes have comparable critical lengths. The plots show that each buckling mode fails to emerge above a certain critical *a* values according to Eq. (9). For each value of *a*, the critical buckling length is plotted for the five lowest buckling modes capable of emerging in the system. The non-linear relationship between the critical buckling length and the buckling mode results in higher buckling modes sometimes emerging at shorter length than lower modes predicting the higher mode is actually the preferred buckling mode. These plots predict kinking as the dominant only for small values of *Ψ*.

The effects of these two dynamics are evident in Fig. 5 where we plot the critical buckling length *N*_*crit*_ as a function of *Ψ* for the first five buckling modes capable of emerging for the given values of *a*. In Fig. 5A and B, for *a* = 1 and *a* = 2all of the five lowest buckling modes are capable of emerging in the system, respectively. Upon increasing to *a* = 5 in Fig. 5C, the first buckling mode is no longer capable of emerging in the system as the critical *a* for the first buckling mode is approximately 3.36. Neither the first nor second mode are present for *a* = 20 in Fig. 5E and subsequent jumps to *a* = 50 and *a* = 100 in Fig. 5F and G, respectively, each result in the loss of additional lower buckling modes.

For the lowest values of *a* = 1, *a* = 2, and *a* = 5 in Fig. 5A–C, we observe the shortest critical buckling length is associated with the lowest present buckling mode and the higher buckling modes emerge at longer lengths in a sequential order. However, we first observe this pattern break down for *a* = 10 in Fig. 5D, where we see the third buckling mode emerge at a much shorter length than the second mode. Interestingly, in this case the second mode also emerges at longer lengths than the fourth, fifth, and sixth modes. This unexpected behavior is due to the non-linear relationship between the critical buckling length *N*_*crit*_ and the buckling mode *n*, as shown in Eq. (8). This non-linearity results in the same behavior for larger values of *a* as Fig. 5E-G show many higher buckling modes emerging at shorter lengths than lower modes.

### Dependence of instability mode on biophysical conditions

To understand whether buckling or kinking is expected to emerge in the growing bacterial chain, we further compare the critical chain lengths for buckling instability to that for kinking instability. The critical kinking length was found in our previous work [20] to be

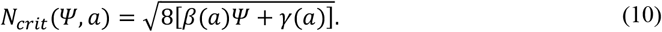

The values of *β* and *γ* are given in Table 2 and the method used to compute them are detailed in the Supplementary Materials.

**Table 2:** Slopes *β* and intercepts *γ* for critical length of kinking emergence. Values are found via linear regression of the results of the perturbation analysis of the rigid-rod model.

| $a$ | $\beta$ | $\gamma$ |
| --- | --- | --- |
| 1 | 3.49 | 0.31 |
| 2 | 3.49 | 0.42 |
| 5 | 3.49 | 0.75 |
| 10 | 3.49 | 1.31 |
| 20 | 3.49 | 2.43 |
| 50 | 3.49 | 5.80 |
| 100 | 3.49 | 11.40 |

Plugging the values from Table 2 to Eq. (10), we plot the critical kinking length as a function of *Ψ* for each specific value of *a* in Fig. 5. For most values of *a*, critical kinking length is longer than the critical buckling length for all or most of the first five buckling modes, suggesting buckling is the dominant mechanical instability for most values of *Ψ* and *a*. This pattern is most prominent for smaller values of *a* as in Fig. 5A for *a* = 1, where the critical buckling length is shorter than the critical kinking for almost any value of *Ψ*. For larger values of *a*, such as *a* = 50 and *a* = 100 in Fig. 5F and G, we observe certain ranges of *Ψ* for which the critical kinking length is shorter than the critical buckling length, and therefore kinking is the dominant instability. These regions correspond to low values of *Ψ*. For comparison with the simulation results, we summarize these findings in Fig. 6A for the specific values of *Ψ* and *a* considered in the simulations in Fig. 3. For each pair of *Ψ* and *a* values, we determine the instability mode, either kinking or a specific buckling mode, that emerges at the shortest critical chain length. There is a clear, kinking dominant region for small *Ψ* and large *a*, while buckling dominates the rest of the physical phase space.

**Figure 6:**
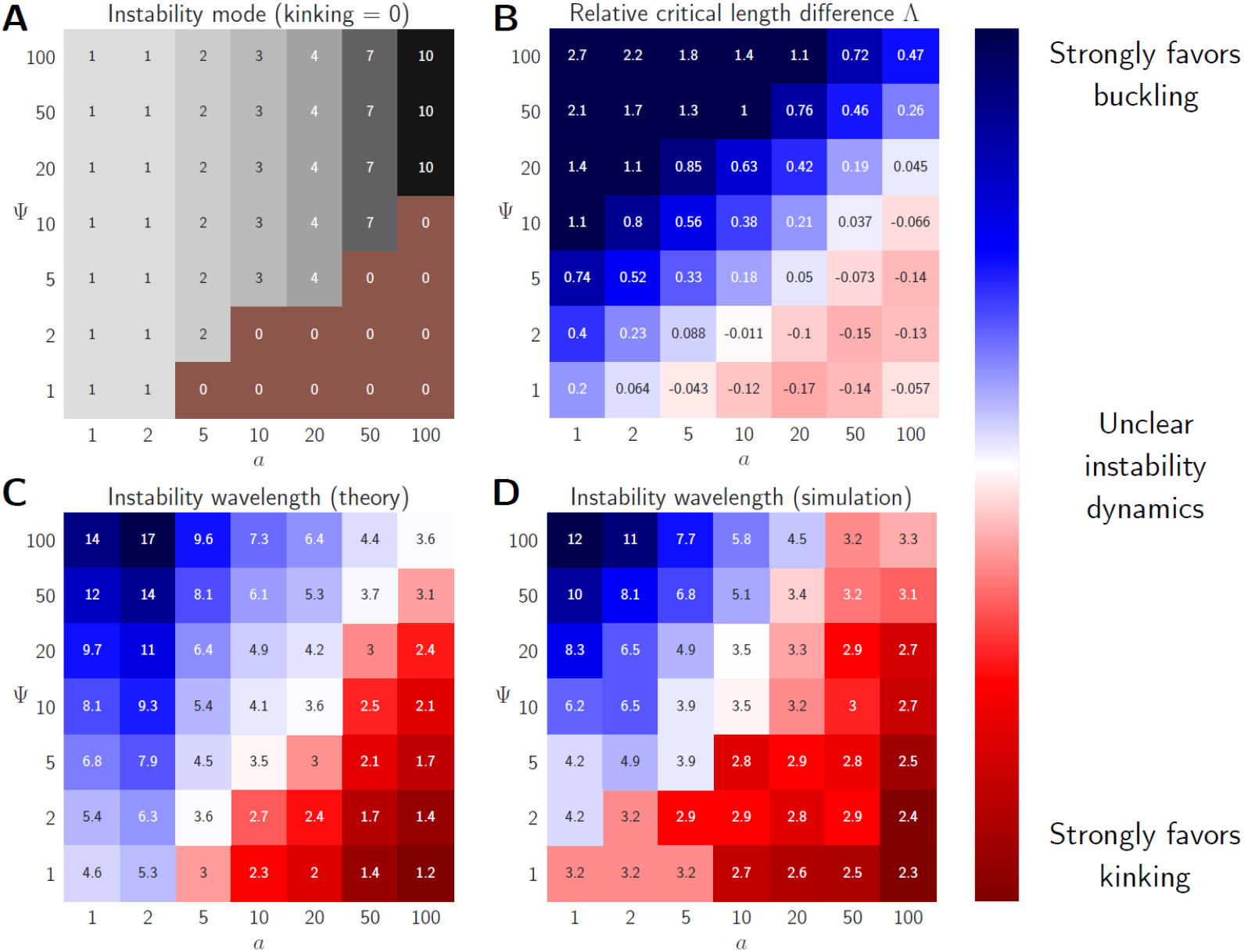
Predicting the instability dynamics from mathematical theory and computer simulations. (A) The predicted instability mode for various points in the *Ψ, a* phase space based on the mode with the shortest critical length from Eqs. (8) and (10). The instability mode *n* = 0 corresponds to kinking, while *n* > 0 corresponds to the *n*-th buckling mode. (B) The relative critical length difference *Λ* for various values of *Ψ* and *a. Λ* measures the difference between the critical kinking length and the shortest critical buckling length relative to critical length of the dominant dynamic. *Λ* > 0 corresponds to buckling as the favored dynamics while *Λ* < 0 corresponds to kinking. (C) The instability wavelength is computed using Eq. (12) from the theoretical critical buckling lengths found in Eq. (8). The wavelength can be interpreted as the number of cells between significant changes in curvature throughout the chain. (D) The instability wavelength computed using Eq. (12) from the simulations in Fig. 3B. The calculations use the curvature profile from the final time step of the simulations at *t* = 100 min.

In Fig. 5, we observe many cases, typically for small values of *Ψ*, where the critical buckling length and critical kinking lengths are very close, suggesting that in these cases it is unclear which instability dynamic is dominant. To understand how broad this region of ambiguity is, we consider the relative critical length difference, a function of *Ψ* and *a*, that compares the critical kinking length with the critical buckling length of the most favorable buckling mode (i.e., the mode with the shortest critical length). The relative critical length difference, denoted by *Λ*, measures the difference between the two predicted critical lengths relative to the critical length of the dominant dynamic (i.e., the instability mode with the shortest critical length):

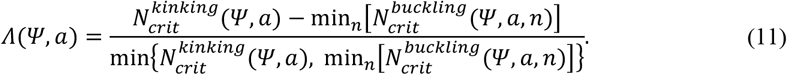

*Λ* is positive if buckling is favored and negative if kinking is favored. Fig. 6B shows the relative critical length difference for the same values of *Ψ* and *a* considered in Fig. 3B and the calculation of this metric is based on the predictions provided in Fig. 5. Using *Λ* as a metric for the instability mode, buckling remains favored for large *Ψ* and small *a* in Fig. 6B, qualitatively agreeing with the predictions from Fig. 6A. However, there is clearly a significant region in the *Ψ, a* phase space in which the critical kinking and critical buckling lengths are comparable and it is unclear which dynamic is favored. Additionally, the relative critical length difference predicts kinking is only slightly favored for small *Ψ* and large *a*, a region in which simulations suggests strongly favors kinking (Fig. 3B).

To quantify the dynamics in the region where the critical kinking and buckling lengths are comparable, we introduce an instability wavelength that characterizes the chain. We apply this metric to the buckling predictions in Eq. (8), assuming the shape of the curvature profile perfectly follows that of the most preferable buckling mode. We compute the instability wavelength as the predicted critical buckling length divided by the associated buckling mode number:

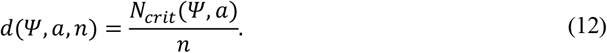

The instability wavelength is interpreted as the distance between points of zero curvature in the chain, assuming the curvature profile shape exactly matches that of the favored buckling mode. *d* is a dimensionless wavelength and therefore quantifies the typical number of cells present between significant changes in curvature. If the cell chain truly exhibits a buckling pattern, the distance between significant changes in curvature would be large relative to the length of a single cell, i.e., *d* ≫ 1. We know the local curvature in adjacent cell linkages will change sharply in kinked chains hence we expect kinking to be dominant if *d* ≲ 1, even if the predicted critical buckling length is shorter than or similar to the critical kinking length. Therefore, the instability wavelength can identify regions in the *Ψ, a* phase space where our models predict buckling emergence as favorable but at such a critical length that the local curvature is not persistent beyond a single cell length. Fig. 6C shows the instability wavelength for the most favorable buckling modes across the *Ψ, a* phase space according to the predictions from Eq. (8). It is helpful to compare these results with those of Figs. 6A and B, where buckling is predicted as the dominant dynamic for large portions of the *Ψ, a*. phase space. It is apparent the instability wavelength provides the desired nuance lacking in the previous predictions. The extremes of Fig. 6C agree with the initial observations from Fig. 3 that buckling is dominant for large *Ψ* and small *a* while kinking prevails for small *Ψ* and large *a*. However, we learn from the instability wavelength that the kinking dominant region is much larger than the predictions in Figs. 6A and B suggest. There is still a significant region for which it remains unclear whether kinking or buckling is the favored dynamic, an unsurprising result since the transition between the two dynamics gradual as shown by the simulations in Fig. 3B.

For additional comparison, we compute the instability wavelengths from the curvature profile at *t* = 100 min in each of the rigid-rod model simulations from Fig. 3. The curvature profile in each of these simulations is computed using the Menger curvature method. This method estimates the curvature in the *i*-th cell-cell linkage using the spatial coordinates of the (*i–*1)-th, *i*-th, and (*i*+1)-th cell linkages. In the analysis of the curvature profile, we count the number of extrema and compute the wavelength as the non-dimensional chain length divided by the number of counted extrema. This computation is fairly analogous to the instability wavelength defined in Eq. (12) and allows us to use a single metric to compare the instability dynamics predicted by the rigid-rod model simulations with those predictions from the theoretical frameworks. The instability wavelengths computed from the simulation data are provided in the Fig. 6D and largely agree with the instability predictions in Fig. 6C.

In sum, we have used several mathematical analyses and metrics to quantify the possibility of kinking versus buckling emergence across the *Ψ, a* phase space. Our mathematical predictions of the critical buckling and kinking lengths provide insight into which dynamic is favored as function of *Ψ* and *a*. We learn buckling is the favored dynamic for large values of *Ψ* and small values of *a*, while kinking dominates for small *Ψ* and large *a*. To clarify the predicted dynamics in between these two extremes of the phase space, we apply metrics such as the relative critical length difference and the instability wavelength. Together these predictions provide a picture of the landscape of mechanical instability emergence across the *Ψ, a* phase space which defines the possible physical regimes in which the bacteria utilizing chain-mediated motility live. In the following section we will apply these predictions along with additional analysis to investigate the biological functions and impacts of mechanical instability emergence in growing bacterial chains.

### Buckling delays chain breakage but reduces sliding efficiency

In the previous sections we examined how the physical conditions of the bacterial chain and its environment dictated the instability mode emergence. We now look for potential biological roles mechanical instability emergence may play in bacterial spreading utilizing chain-mediated motility.

It has been shown that stress in a perfectly straight chains follows a parabolic profile [20, 25, 29] and increases exponentially at a rate twice that of the cell growth rate [20]. This exponential increase in stress is unsustainable for long periods of time and one would expect the growing to stress to eventually result in mechanical failure. Perturbation analysis of the rigid-rod model for growing bacteria chains predicts a critical stress that depends on the non-dimensional parameters *Ψ* and *a* [20]. This critical stress is the same as the critical kinking stress associated with the critical kinking length. However, we can also interpret this critical kinking stress as the critical stress associated with breakage of the cell-cell linkages since sharp kinking is likely analogous to breakage of the linkages. Given that the stress in the perfectly straight growing cell chain increases exponentially in time, it would be advantageous for the bacterial chain to develop additional mechanisms that assist the chain in avoiding breakage. Buckling may be one such mechanism.

The simulations in Fig. 3B consist of bacterial chains initialized with all cells laying perfectly horizontal along the *x*-axis except for the two centermost cells. The centermost cell-cell linkage of this perfectly straight chain has been perturbed so that the angle between the centermost cells is *Δθ*. This small perturbation just barely pushes the chain system away form the perfectly straight state and allows for instability emergence in the system that would theoretically not be possible if the chain was truly perfectly straight. We typically observe the perturbation decrease such that *Δθ* approaches zero during the initial stages of the simulation. As such for a stretch of time during the early portions of the simulations, the chains are nearly perfectly straight with the expected parabolic stress profile that increases exponentially in time (see *t* = 5, 45 min in Fig. 7A and B). The chain continues growing in this nearly straight configuration until a mechanical instability emerges. In many cases, the chain grows straight for a long enough period of time that the largest stress in the chain (the stress in the centermost linkage) exceeds the predicted critical breaking stress. In simulations with this behavior, we expect kinking to emerge as the sharp bending is intuitively comparable to chain breakage.

**Figure 7:**
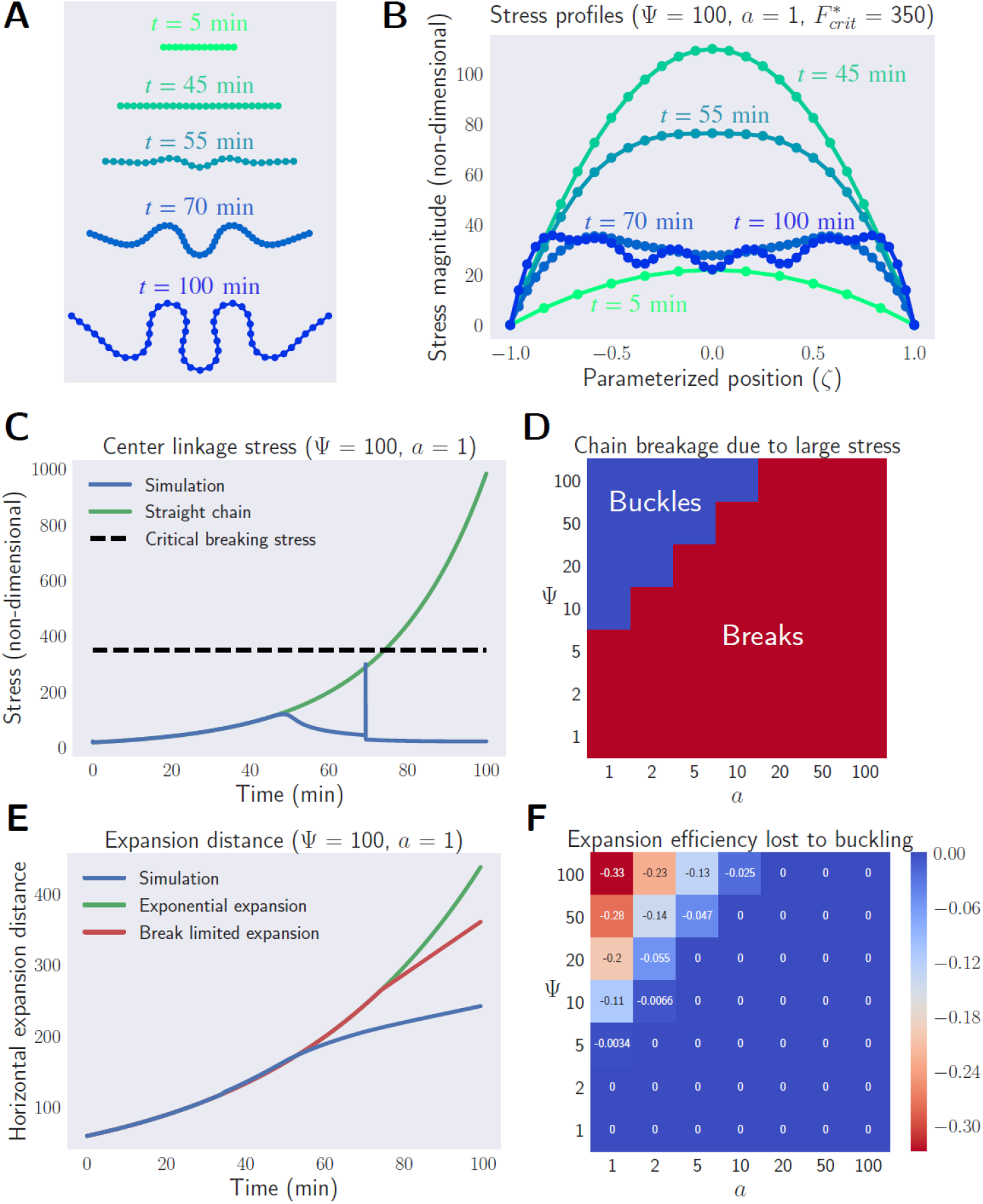
Buckling reduces the stress in growing cell chains at the cost of the expansion efficiency. (A) Simulated chain time series for *Ψ* = 100 and *a* = 1. (B) Non-dimensionalized stress magnitude profile at each time point in (A). (C) Magnitude of the stress in the centermost linkage as a function of time from the simulation in (A). The discontinuity in the stress at *t* ≈ 70 min is a transient artifact of the cell division implementation in the rigid-rod model that likely does not emerge in the real world system and does not affect our conclusions. (D) Simulation results across the *Ψ*–*a* phase space depicting whether the stress in the chain ever exceeds the critical breaking stress within the simulation time (red) or reduces its stress by buckling prior reaching the critical breaking stress (blue). (E) The horizontal expansion distance as a function of time (blue) for the chain simulated in (A) compared with exponential expansion (green) and break limited expansion (red). (F) The loss in horizontal expansion efficiency due to buckling across the *Ψ, a* phase space relative to the break limited expansion. We assume chains that reach critical stress prior to buckling follow the break limited expansion rate.

Interestingly, buckling results in a significant change in the stress profile of the growing chain. Fig. 7A shows a time series of the chain configuration for an initially perturbed chain and Fig. 7B shows the evolution of the magnitude of its stress profile. The chain grows nearly straight and exhibits an increasing parabolic stress profile for the early stages of the simulation. At *t* ≈ 55 min a buckling pattern emerges throughout the chain and, strikingly, we observe a drop in the stress profile. Initially the parabolic stress profile flattens in the center and as the buckling becomes more prominent the stress in the center drops below that of the surrounding linkages. This decrease in the stress profile helps the chain avoid breakage. Fig. 7C shows the stress in the center linkage of the simulated chain as a function of time. The stress initially increases exponentially as we expect for a nearly straight chain, but the stress in the center linkage beings to drop at *t* ≈ 55 min in correspondence with the emergence of the buckling pattern. Fig. 7D presents this dynamic across the *Ψ, a* phase space detailing whether or not the stress in the chains simulated in Fig. 3B ever exceeds the predicted critical breaking stress. We see two clear regions in the *Ψ, a* phase space. For large *Ψ* and small *a*, the chains avoid breakage through the buckling emergence. This region agrees with the buckling dominant regimes identified in Fig. 6. The decrease in stress resulting from buckling emergence allows the chain to continue growing while avoiding stress levels that would cause chain breakage.

While buckling potentially helps chains avoid breakage, it does have a drawback in regard to the horizontal expansion. The perturbed cell chains in Fig. 3B initially grow nearly straight along the horizontal *x*-axis. As buckling emerges in the cell chain, the most curved parts of the chain extend in the *y*-direction. If chain-mediated sliding motility is utilized as a means to expand in a directed, one-dimensional fashion, then buckling would decrease the efficiency of such expansion. In order to quantify this decrease in efficiency, we compare the expansion of buckled chains with the expansion of straight chains. Analysis of a simplified population model for growing straight chains combined with the perturbation analysis of the rigid-rod model showed regular occurrence of breaks in the chains growing along a horizontal axis limited the expansion to a constant rate despite the exponential growth of individual chains [20]. This analysis predicted the break limited expansion rate for straight chains as a function of *Ψ* and *a* to be

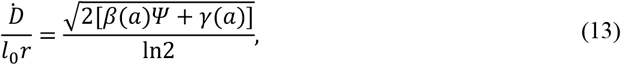

where *β* and *γ* are the same *a*-dependent coefficients presented in Table 2.

Fig. 7E shows the expansion distance for a simulated chain that exhibits a buckling pattern prior to stress in the chain exceeding the critical breaking stress. We compare the expansion of this buckled chain with the expansion of the break limited straight chains. The simulated chain initially expands exponentially until a buckling instability emerges in the system. In comparison, the perfectly straight chain also grows exponentially, in this case until a break occurs. The buckling pattern emerges in the simulated chain at a time such that the stress in the chain does not reach the critical breaking stress, thus allowing the chain to avoid a break. However, the expansion from this point onward is less efficient than the exponential expansion. Meanwhile, the perfectly straight chain expands exponentially until its center stress exceeds the critical breaking stress, from which point onward the expansion is linear. The expansion of the simulated chain that buckles is less efficient than the predicted break limited expansion of the perfectly straight chain. Fig. 7F shows the expansion rates for simulated chains relative to the break limited expansion rate predicted by Eq. (13) for various pairs of *Ψ* and *a* values. If the stress in the simulation ever exceeds the critical breaking stress, we assume that chain experiences a break and its final expansion rate will match the predicted break limited expansion. In the region where buckling is dominant (large *Ψ*, small *a*), the simulations predict less efficient expansion relative to the break limited expansion. However, for the simulations close to the buckling-kinking boundary, the loss of efficiency is much smaller. For these simulations, the critical buckling stress is slightly less than the critical breaking stress such that buckling emerges close to the point in time we predict chain breakage. This allows the chain to maximize the amount of time it expands exponentially before the bucking instability emerges and the expansion diverges to a less efficient rate. These results suggest buckling may be a useful mechanism for avoiding or at least delaying chain breakage at the cost of expansion efficiency. However, the loss of expansion efficiency is less if the critical buckling stress is only slightly smaller than the critical breaking stress.

We have shown growing cell chains will exhibit one of two mechanical instabilities; kinking and buckling. Kinking results in a buildup in stress in the cell-cell linkages and eventually leads to sharp bending in these linkages that corresponds to chain breakage. Buckling results in the emergence of smooth curvature patterns and decreases the stress in the growing chain, a dynamic that allows the chain to avoid or delay a cell-cell linkage failure. Both of these mechanical instabilities prevent the chain from achieving highly efficient exponential expansion.

## Discussion

We consider the emergence of two modes of mechanical instability in growing bacterial cell chains. Using experimental observations and physics-based simulations, we demonstrate bacterial chains exhibit either a buckling instability or a kinking instability depending on the physical properties of the chain and its surrounding environment. The buckling instability results in the emergence of regions of large curvature throughout the chain, while the kinking instability causes sharp, localized bending that can lead to mechanical failure of the cell-cell linkages. We introduce a continuum mechanics framework for growing cell chains and apply this framework in conjunction with the existing rigid-rod model for cell chains. Both models contain the same fundamental physical forces and depend on the same physical parameters making for easy comparison of each model’s predictions. With these models, we quantify the expected instability mode as a function of the physical conditions.

Perturbation analysis of the continuum mechanics framework along with our curvature profile ansatz for the buckling modes (Eq. (7)) results in predictions for the critical lengths associated with the emergence of a particular buckling mode as a function of the physical parameters. We compare these critical buckling lengths with the critical kinking lengths predicted by a similar perturbation analysis of the rigid-rod model to determine the most favorable instability as a function of our physical parameters, *Ψ* and *a*. This analysis reveals buckling is the expected dominant instability dynamic if the cell-cell linkages are strong relative to the viscous drag of the environment (large *Ψ*, Eq. (1)) and if the anisotropic drag ratio is small (small *a*, Eq. (2)).Conversely, kinking is the expected dominant instability dynamics if the linkages are weak relative to the environmental drag (small *Ψ*) and if the anisotropic drag coefficient is large (large *a*). These predictions uncover an interesting non-linear relationship between the critical buckling length and the given buckling modes allowing for higher buckling modes to potentially emerge at shorter lengths than the lower mode given the proper physical conditions. We demonstrate the existence of a critical value of the anisotropic drag coefficient *a* for each buckling mode above which emergence of that specific buckling mode is no longer possible. We apply additional analysis and introduce metrics such as the relative critical length difference and the wavelength to further characterize the instability dynamics in between the extremes of the physical phase space. These metrics reveal a region of the phase space in which the exact instability mode is unclear and details the transition into the respective regions where buckling and kinking are the dominant dynamic.

Using the physics-based simulations from the rigid rod, we consider the biological impacts of buckling and kinking. Simulations of buckled chains reveal that the emergence of buckling reduces stress in the chain making its cell-cell linkages less prone to breakage. However, this reduction in stress is coupled with a decrease in the efficiency of the chain’s horizontal expansion from the exponential expansion of perfectly straight chains. This presents a trade-off between buildup in mechanical stress and high-speed one-dimensional expansion. The loss of expansion efficiency due to buckling emergence is often more significant than the loss of efficiency due to regular chain breakages, but if the critical buckling stress threshold is only slightly lower than the threshold for breaking, the loss of efficiency is comparable. It is also important to recognize the predictions with regard to horizontal expansion with the occurrence of regular breaks is a best-case scenario–in reality, repeated breakages may actually lead to less efficient expansion than is predicted and it is possible that buckling limited expansion is more effective than break limited expansion under certain conditions. In general, less efficient expansion is unavoidable for growing cells chains–exponential expansion is known to cause an exponential increase in stress that is not sustainable. The analysis of mechanical stress and horizontal expansion in buckled chains reveals that buckling may serve as a mechanism that allows bacterial chains to delay or even avoid breakage of its cell-cell linkage at the cost of less efficient expansion. Whether expansion is limited by buckling emergence or linkage failure, as well as the exact implication of these behaviors on horizontal expansion, depend on the physical conditions of the chain and its surrounding environment.

Our mathematical models reveal that the instability dynamics for growing cell chains depend on only two free parameters: *Ψ* (Eq. (1)) and *a* (Eq. (2)). It is easy to interpret the results with respect to *a* since it is simply the anisotropic ratio between the parallel and perpendicular drag coefficients (with respect to the cell body). However, since *Ψ* contains four different physical parameters, its interpretation is more complex. We have qualitatively described *Ψ* as the ratio between the cell-cell linkage strengths (*m* in the continuum model, *k*_*b*_ in the rigid rod model) and the viscosity of the substrate (*μ* in both models). Examining the ratio between these two properties is straightforward. For example, it is reasonable that it is necessary to increase mechanical linkage strength in a more viscous environment in order to achieve the same dynamics as in a less viscous environment. Variations in each of these two physical parameters individually are also fairly intuitive. It is reasonable to expect a chain to be more susceptible to localized bending if the linkages are relatively weak or if the viscosity is relatively high. However, the exact mechanical implications of variations in the cell length (*l*_0_) or the cell growth rate *rr* are not as clear. Non-dimensionalization of both the continuum mechanics and rigid rod frameworks simplifies these interpretations. Since all predictions are in terms of the two non-dimensional parameters *Ψ* and *a*, they provide predictions for variations in each individual parameter. This also makes experimental testing of these predictions straightforward as it is possible to isolate the predictions in terms of which ever physical parameter that is isolated as the independent variable for a set of experiments.

Our results highlight the rich mechanical dynamics of growing bacteria chains and provide useful information for understanding the broader behavior of bacterial populations. The buckling and kinking instability dynamics are directly related to the mechanical efficacy of individual chains as well as the effectiveness of population expansion. Understanding these mechanical dynamics is crucial as bacterial chaining is likely an important dynamic in many aspects of the bacterial lifecycle. The formation of bacterial chains plays a role in the early formation of biofilms [30-33] and recent studies identify forced bacteria chaining as an effective immune response against infection [34, 35]. Given these connections to multiple aspects of the bacterial lifecycle, the mechanical insights to the chain dynamics from our mathematical models may lead to a better understanding of biofilm formation and pathogenicity in many bacterial species. Continued investigation of these mechanical dynamics is crucial to fully understanding the roles and impacts of chain formation in the life-cycle of many bacteria.

The continuum and rigid rod modeling frameworks are both generic and therefore can be applied to any bacteria that utilizes chain-mediated sliding motility. It is important to note the differences between these two models in order to apply them properly. The continuum mechanics framework assumes the chain is uniform and therefore does not include discrete weak point. This framework provides continuous functions for physical quantities and therefore was the best choice for analyzing global curvature dynamics in our buckling emergence investigation. The rigid-rod model assumes discrete cell bodies that are rigid and inflexible. In reality cell bodies flexible to some degree so investigations where the flexibility of individual cells is important may prefer the continuum mechanics model. While the rigid rod assumption is less realistic, it does allow for the implementation of discrete cell-cell linkages, an important feature when consider mechanical efficacy of growing cell chain. The implementation of the discrete weak point is more realistic for modeling chains where the peptidoglycan cell walls are much stiffer than the walls maintaining the cell-cell linkages. The rigid-rod model also offers the advantage of including a system of ordinary differential equations that can be used to simulate the dynamics of the growing chain.

The linear stability analysis of the continuum mechanics model used to derive the critical buckling conditions is essentially an eigenvalue problem that we solve numerically. We define an ansatz for the curvature profiles and use this function to numerically test if the system is stable or unstable against perturbation to the curvature as function of the chain length. However, the ansatz we introduce in Eq. (7) is not an exact solution to this eigenvalue problem, but rather an approximate solution. Solving for the exact solution requires one can ensure Eq. (6) is negative definite, i.e., negative for any curvature profile. Since we cannot test all possible curvature profile functions, we make the numerical ansatz based on the curvature profiles we observe in the rigid rod simulations. This ansatz turns the eigenvalue problem into simple algebra, without sacrificing much insight into the mechanical dynamics of buckling emergence and its biological impacts. This method of decomposing the perturbation into a series of modes is a classical method for linear stability analysis and has been used to examine the emergence of Turning patterns in reaction-diffusion systems [36].

Overall, our investigation provides new insights to the mechanical instability dynamics that emerge in chain-mediated sliding motility. Our experimental and simulation observations show two different mechanical instabilities, buckling and kinking, that emerge in growing bacterial cell chain. The two instabilities are associated with different mechanical outcomes: bucking results in smooth curvature emergence leading to the chain folding on itself; kinking causing sharp, localized bending that leads to breakage of cell-cell linkages. Our models and analysis highlight the key physical conditions governing instability emergence, including cell body length, cell body growth rate, cell-cell linkage strength, substrate viscosity, and anisotropic drag strength. Our predictions regarding instability emergence are important in developing a deeper understanding of chain-mediated sliding motility, a likely widespread mechanism offering the potential for extremely fast population expansion. Given the ability of many pathogenic bacteria to form chains and the known connections between chain formation and immunity resistance, it is likely further investigation into the mechanical dynamics of growing bacterial chain will provide valuable insights to the pathogenic dynamics of many bacteria.

## Acknowledgements

This work was supported by NIH grant 1R35GM138370 to J.C.

