## Supplementary Materials for "Kinking and buckling instability in growing bacterial chains"

#### A Continuum mechanics model for growing cell chains

##### A.1 Connection to Euler's column – Overview

In Chapter 2, there are angular springs which favor linear alignment of the cells in the chain. In the macroscopic model, the chain is elastic, with resistance to bending. Let's suppose a section of rod is subject to compressive pressure stresses acting across its endpoints. As an analogy, imagine squeezing a straw between your thumb and fore-finger. Squeeze lightly, and the straw remains straight. Squeeze hard enough, and the straw bows into an arc. This is the phenomenon of buckling, first examined by Euler in his analysis of a loaded column. Our analysis of a buckling bacteria chain is to be based on the mechanics of the chain bending in response to drag induced forces along its length. The first order of business is to formulate this one-dimensional continuum mechanics.

##### A.2 Chain continuum mechanics

We begin with descriptions of *internal* forces representing interactions between material elements of the chain with each other. There are forces of resistance against bending, and contact forces of a rigid cell pushing against its neighbors. The descriptions of these forces dont involve time evolution, and it is convenient to employ a parametric representation $\mathbf{x}(s)$ of the chain curve with respect to arclength $s$, $0 < s < L$. In Fig. 3.8, $\mathbf{t}(s)$ denotes the unit tangent oriented in the direction of increasing $s$, and $\mathbf{n}(s)$, the unit normal counterclockwise with respect to $\mathbf{t}(s)$. The angle between the tangent and the horizontal is denoted by $\theta(s)$.

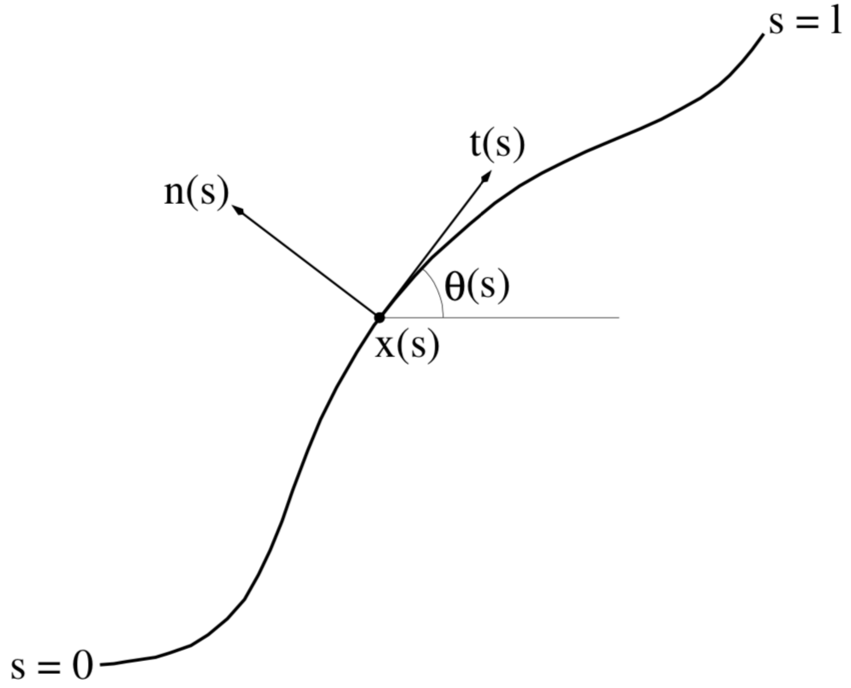


**Figure 3.8: Parametric representation of the continuous cell chain.**

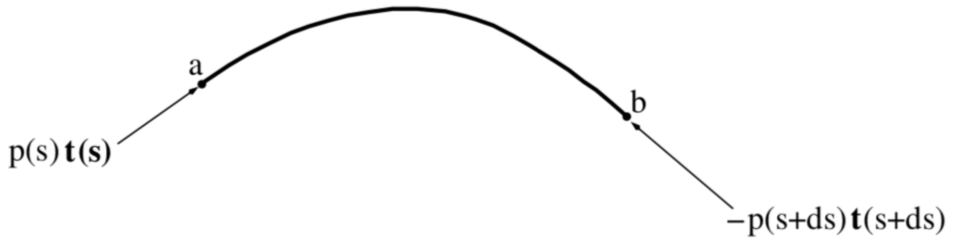


**Figure 3.9: Visualization of the action of contact forces on segment of the chain.**

Fig. 3.9 visualizes the action of contact forces on the segment of chain in the arclength interval $(s,\ s+ds)$. The cells to the left of point a exert a push $p(s)\mathbf{t}(s)$ on $ab$. Think of a cell as a very stiff spring: When compressed it exerts opposite and equal forces on adjacent cells to the left and right, in line with itself. The magnitude of this force is called pressure. In Fig. 3.9, $p(s)$ is the pressure of the push from the left acting across $a$, and $p(s+ds)$, the pressure of the push from the right acting across $b$. The net contact force on $ab$ is

$$-p(s+ds)\mathbf{t}(s+ds)+p(s)\mathbf{t}(s)\sim-\partial_s(p\mathbf{t})ds, \tag{3.17}$$

and we discern the contact force per unit length

$$-\partial_s(p\mathbf{t}). \tag{3.18}$$

We discern bending forces from the work required to vary the *bending energy*

$$U=\frac{m}{2}\int_0^L(\partial_s\theta)^2\,ds. \tag{3.19}$$

The essential idea is the principle of virtual work. Here is a brief review in the simplest context of n discrete degrees of freedom: Let $U(q_1,...,q_n)$ be the potential energy as a function of the coordinates $q_1,...,q_n$. The generalized force acting on the $i$-th degree of freedom is

$$\phi_i=-\partial_i U. \tag{3.20}$$

The work required to impose incremental changes $\delta q_1,...,\delta q_n$ is

$$-\sum_i\phi_i\delta q_i. \tag{3.21}$$

Notice the minus sign: *positive* work when you move against the force. Substituting Eq. 3.20 for $\phi_i$, we have

$$-\sum_i\phi_i\delta q_i=\sum_i\partial_i U\delta q_i=\delta U \tag{3.22}$$

In summary: $\delta U$ is a linear combination of the $\delta q_i$. The coefficients of this linear combination are $\phi_i$, the negatives of the generalized forces.

The bending forces per unit length follow from a continuum version of the same analysis. We seek the force per unit length $\phi=\phi_1\mathbf{e}_1+\phi_2\mathbf{e}_2$ so that

$$\delta U=-\int_0^L\{\phi_1\delta x_1+\phi_2\delta x_2\}\,ds. \tag{3.23}$$

Its variation of bending energy Eq. 3.19 due to deforming the curve is

$$\begin{aligned}\delta U=&m\int_0^L\partial_s\theta\partial_s\delta\theta ds=\\ &m\left[\partial_s\theta\delta\theta\right]_0^L-m\int_0^L\partial_{ss}\theta\delta\theta ds.\end{aligned} \tag{3.24}$$

Here $m$ is the bending modulus. The angle $\theta(s)$ is related to the components of $\mathbf{x}(s)$ by

$$\theta = \arctan \frac{t_2}{t_1}. \tag{3.25}$$

Here $t_1 := \partial_s x_1, t_2 := \partial_s x_2$ are the components of the tangent vector. From Eq. 3.25 we find

$$\delta\theta = t_1 \partial_s \delta x_2 - t_2 \partial_s \delta x_1. \tag{3.26}$$

Substitution into Eq. 3.24 gives

$$\begin{aligned} \delta U =& m \left[\partial_s \theta \delta\theta\right]_0^L - m \int_0^L \partial_{ss}\theta \left(t_1 \partial_s \delta x_2 - t_2 \partial_s \delta x_1\right) ds = \\ & \left[\partial_s \theta \delta\theta - m \partial_{ss}\theta t_1 \delta x_2 + m \partial_{ss}\theta t_2 \delta x_1\right]_0^L + \\ & m \int_0^L \left\{\partial_s \left(\partial_{ss}\theta t_1\right) \delta x_2 - \partial_s \left(\partial_{ss}\theta t_2\right) \delta x_1\right\} ds. \end{aligned} \tag{3.27}$$

Comparing Eqs. 3.22 and 3.27, we discern

$$\phi_1 = m \partial_s \left(\partial_{ss}\theta t_2\right), \phi_2 = -m \partial_s \left(\partial_{ss}\theta t_1\right),$$

and the force per unit length associated with bending is

$$\phi = -m \partial_s \left(\partial_{ss}\theta \left(-t_2 \mathbf{e}_1 + t_1 \mathbf{e}_2\right)\right) = -m \partial_s \left(\partial_{ss}\theta \mathbf{n}\right) \tag{3.28}$$

The sum of forces per unit length due to contact and bending is, from 3.18 and 3.28

$$-m \partial_s \left(\partial_{ss}\theta \mathbf{n} + p \mathbf{t}\right). \tag{3.29}$$

For chains with free ends, we have the natural boundary conditions, that $\partial_s \theta$ and $\partial_{ss}\theta$ vanish at $s = 0, L$. There is no compression at free ends, so the pressure $p$ vanishes at the ends as well.

The drag forces per unit length on the moving, growing chain balance the mechanical force per unit length Eq 3.29. What are they? It is convenient to change the parametric representation of the chain curve to $\mathbf{x}(\zeta, t)$, where the curve parameter $\zeta$ ranges between $-1$ and 1 (like in section I). The trajectory $\mathbf{x} = \mathbf{x}(\zeta, t)$ with $\zeta$ fixed "follows" cells, in that the velocity of a cell at $\mathbf{x}(\zeta, t)$ is $\partial_t \mathbf{x}(\zeta, t)$. The arclength between $\zeta_1$ and $\zeta_2$ is

$$\frac{L(t)}{2} \left|\zeta_2 - \zeta_1\right| \tag{3.30}$$

which grows for increasing $L(t)$. Hence, the conversion between $s$ and $\zeta$ derivatives is

$$\partial_s = \frac{2}{L} \partial_\zeta. \tag{3.31}$$

In the $\zeta$-parametrization, the drag force per unit length may be expressed as

$$\mathbf{f} = -\mu\left(\partial_t \mathbf{x}\cdot\mathbf{t}\right)\mathbf{t} - \nu\left(\partial_t \mathbf{x}\cdot\mathbf{n}\right)\mathbf{n} \tag{3.32}$$

where $\mu$ and $\nu$ are longitudinal and lateral drag coefficients. The dynamical equation which results from the balance of drag and mechanical forces is

$$\mathbf{f} = \left(\frac{2}{L}\right)^3 m\partial_\zeta\left(\partial_{\zeta\zeta}\theta\mathbf{n}\right) + \frac{2}{L}\partial_\zeta(p\mathbf{t}) \tag{3.33}$$

We used Eq. 3.31 to convert $s$-derivatives in Eq. 3.29 into $\zeta$-derivatives in Eq. 3.33.

#### A.3 Linearized dynamics about a straight line chain

There is the straight line chain solution of Eq. 3.33, with

$$\mathbf{x}(\zeta,t) = \frac{L(t)}{2}\zeta\mathbf{e}_1. \tag{3.34}$$

For this chain, $\theta(\zeta,t)\equiv 0$ and $\mathbf{t}\equiv\mathbf{e}_1$. We linearize the dynamics of Eq. 3.33 about this line chain solution. The gauge parameter $\epsilon$ measures the deviation from the straight line solution. The expansions of $\mathbf{x}(\zeta,t), p(\zeta,t)$ in the limit $\epsilon\to 0$ are

$$\mathbf{x}(\zeta,t) = \frac{L(t)}{2}\zeta\mathbf{e}_1 + \epsilon Y(\zeta,t)\mathbf{e}_2 + O\left(\epsilon^2\right), \tag{3.35}$$

$$p(\zeta,t) = P(\zeta,t) + O\left(\epsilon^2\right). \tag{3.36}$$

There are no $O(\epsilon)$ components of $x_1(\zeta,t)$ and $p(\zeta,t)$. Suppose we included them. We find that the $O(\epsilon)$ term of $x_1(\zeta,t)$ is a function of time only, independent of $\zeta$. The 1-velocity would have an $O(\epsilon)$ uniform translation component and the net drag force on the entire chain would not vanish. With no $O(\epsilon)$ term of $\partial_t x_1(\zeta,t)$, the self-consistent boundary value problem for the pressure shows that it has no $O(\epsilon)$ term.

First, look at the $\epsilon\to 0$ limits of tangent and normal $\mathbf{t},\mathbf{n}$ and the angle $\theta$. The expansion of the tangent vector is

$$\mathbf{t} = \frac{2}{L}\partial_\zeta\mathbf{x} = \mathbf{e}_1 + \frac{2\epsilon}{L}\partial_\zeta Y\mathbf{e}_2 + O\left(\epsilon^2\right). \tag{3.37}$$

The expansion of the normal is

$$\mathbf{n} = \mathbf{e}_2 - \frac{2\epsilon}{L}\partial_\zeta Y\mathbf{e}_1 + O\left(\epsilon^2\right). \tag{3.38}$$

From (38) we see that the expansion of the angle $\theta$ is

$$\theta = \frac{2\epsilon}{L}\partial_\zeta Y + O\left(\epsilon^2\right). \tag{3.39}$$

The expansion of the velocity $\partial_t \mathbf{x}$ is

$$\partial_t \mathbf{x} = \frac{\dot{L}}{2}\zeta \mathbf{e}_1 + \epsilon \partial_t Y \mathbf{e}_2 + O\left(\epsilon^2\right) \tag{3.40}$$

From Eqs. 3.37, 3.38, and 3.40, we find that the components of drag force per unit length in Eq. 3.32 have expansions,

$$f_1 = -\mu \frac{\dot{L}}{2}\zeta + O\left(\epsilon^2\right), \tag{3.41}$$

$$f_2 = -\epsilon\nu\partial_t Y - \epsilon(\mu - \nu)\frac{\dot{L}}{L}\zeta\partial_\zeta Y + O\left(\epsilon^2\right). \tag{3.42}$$

Similarly, we expand the the RHS of Eq. 3.33. Equating its 1 and 2 components to $f_1, f_2$ according to Eqs. 3.41 and 3.42, we find leading order equations for $Y$ and $P$

$$-\mu\frac{\dot{L}}{2}\zeta = \frac{2}{L}\partial_\zeta P \tag{3.43}$$

$$-\nu\partial_t Y + (\nu - \mu)\frac{\dot{L}}{L}\zeta\partial_\zeta Y = \left(\frac{2}{L}\right)^4 m\partial_{\zeta\zeta\zeta\zeta} Y + \left(\frac{2}{L}\right)^2 \partial_\zeta\left(P\partial_\zeta Y\right). \tag{3.44}$$

The solution of Eq. 3.43 for $P$ subject to zero boundary conditions at $\zeta = -1, 1$ reproduces the pressure field in Eq. **??** of the straight line chain. Given $P$, the PDE Eq. 3.44 becomes

$$-\nu\partial_t Y - (\mu - \nu)\frac{\dot{L}}{L}\zeta\partial_\zeta Y = \left(\frac{2}{L}\right)^4 m\partial_{\zeta\zeta\zeta\zeta} Y + \frac{\mu}{2}\frac{\dot{L}}{L}\partial_\zeta\left(\left(1-\zeta^2\right)\partial_\zeta Y\right) \tag{3.45}$$

The linearizations of the boundary conditions, that $\partial_\zeta\theta, \partial_{\zeta\zeta}\theta$ vanish at the ends are

$$\partial_{\zeta\zeta} Y = 0, \partial_{\zeta\zeta\zeta} Y = 0 \tag{3.46}$$

at $\zeta = -1, 1$

#### A.4 Buckling

We analyze the linearized stability against deformations of the chain from a straight line. Specifically, do perturbations of curvature grow or decay? Under the linearized limit process, the representation of curvature is

$$\partial_s\theta = \frac{2}{L}\partial_\zeta\left(\epsilon\frac{2}{L}\partial_\zeta Y\right) + O\left(\epsilon^2\right),$$

or

$$\partial_s\theta = \epsilon\kappa + O\left(\epsilon^2\right), \quad \kappa := \left(\frac{2}{L}\right)^2 \partial_{\zeta\zeta} Y. \tag{3.47}$$

We take

$$\frac{1}{2}\int_{-1}^{1}\kappa^2 d\zeta \tag{3.48}$$

as a metric of the curvature profile. The growth or decay of curvature is informed by an integral identity that derives from Eqs. 3.45 and 3.46. In place of $Y$, introduce the state variable $Z$ so

$$Y = \left(\frac{L}{2}\right)^2 Z \tag{3.49}$$

We find that

$$\kappa = \partial_{\zeta\zeta} Z. \tag{3.50}$$

The boundary value problem for $Z$ is

$$\begin{gathered} -\nu\partial_t Z = \left(\frac{2}{L}\right)^4 m\partial_{\zeta\zeta\zeta\zeta} Z + \\ \mu\frac{\dot{L}}{L}\left\{\zeta\partial_\zeta Z + \frac{1}{2}\partial_\zeta\left(\left(1-\zeta^2\right)\partial_\zeta Z\right)\right\} + \\ \nu\frac{\dot{L}}{L}\left\{2Z - \zeta\partial_\zeta Z\right\}, \end{gathered} \tag{3.51}$$

in $-1 < \zeta < 1$, subject to

$$\partial_{\zeta\zeta} Z = 0, \partial_{\zeta\zeta\zeta} Z = 0 \tag{3.52}$$

at $\zeta = -1, 1$. Differentiating Eq. 3.51 with respect to $\zeta$ twice and evoking Eq. 3.50,

$$-\nu\partial_t\kappa = \left(\frac{2}{L}\right)^4 m\partial_{\zeta\zeta\zeta\zeta}\kappa + \mu\frac{\dot{L}}{2L}\partial_{\zeta\zeta}\left(\left(1-\zeta^2\right)\kappa\right) - \nu\frac{\dot{L}}{L}\zeta\partial_\zeta\kappa \tag{3.53}$$

in $-1 < \zeta < 1$. The boundary conditions on $\kappa$ are

$$\kappa = 0, \partial_\zeta\kappa = 0 \tag{3.54}$$

at $\zeta = -1, 1$. Multiply Eq. 3.53 by $\kappa$ and integrate over $-1 < \zeta < 1$. After integrations by parts and use of the boundary conditions Eq. 3.54 we arrive at

$$-\frac{\nu}{2}\frac{d}{dt}\int_{-1}^{1}\kappa^2 d\zeta = \left(\frac{2}{L}\right)^4 mQ[\kappa] \tag{3.55}$$

where $Q[\kappa]$ is the quadratic form

$$Q := \int_{-1}^{1}\left\{\left(\partial_{\zeta\zeta}\kappa\right)^2 - \lambda(t)\left(1-\zeta^2\right)\left(\partial_\zeta\kappa\right)^2 + \frac{\nu-\mu}{\mu}\lambda(t)\kappa^2\right\} d\zeta \tag{3.56}$$

Here, $\lambda(t)$ is the function of time

$$\lambda := \frac{1}{2^5}\frac{\mu}{m}L^3\dot{L}. \tag{3.57}$$

If the quadratic form is positive definite, then $\frac{1}{2}\int_{-1}^{1}\kappa^2 d\zeta$ is decreasing whenever $\kappa$ is not identically zero. This indicates that a straight line chain is stable. The existence of $\kappa(\zeta)$ with $Q[\kappa] < 0$ indicates loss of stability. We demonstrate the existence of a critical value $\lambda_* > 0$ so $\lambda(t) < \lambda_*$ indicates stability, and $\lambda(t) > \lambda_*$ indicates instability.

For an exponentially growing chain with $l \propto e^{rt}$, we have $\dot{L} = Lr$, and then

$$\lambda(t) = \frac{1}{2^5}\frac{\mu r}{m}L^4(t). \tag{3.58}$$

Apparently, the chain starts buckling when

$$L > 2\,(2\lambda_*)^{\frac{1}{4}}\left(\frac{m}{\mu r}\right)^{\frac{1}{4}} \tag{3.59}$$

Let $l_0$ be the length of a cell just after it divides. Then $N := \frac{l}{l_0}$ is the number of cells just after a division. We may reformulate Eq. 3.59 as

$$N > 2\,(2\lambda_*)^{\frac{1}{4}}\left(\frac{m}{\mu l_0^4 r}\right)^{\frac{1}{4}} \tag{3.60}$$

The dimensionless parameter

$$:= \frac{m}{\mu l_0^4 r} \tag{3.61}$$

is similar to $\Psi$ in [47]. In place of the bending modulus $m$, they have the angular spring constant $k_b$. The continuum limit of their discrete bending energy yields the identification $m = k_b l_0$, so $\Psi$ in Eq. 3.61 reduces to $\Psi$ in [47].

### B Numerically deriving the buckling condition

Appendix 3.A introduces the continuum mechanics framework for analyzing the dynamics of growing cell chains. Linearization of this model demonstrates the existence of a critical buckling length associated with the buckling emergence. The linearization and following analysis yields Eq. 3.56 which essentially defines an eigenvalue problem where the eigenvalues $\lambda_*$ provide the critical buckling lengths and the eigenfunctions $\kappa(\zeta)$ define the respective buckling mode curvature profiles. We solve this stability problem numerically by making an ansatz for the eigenfunctions. This ansatz turns the eigenvalue problem into straightforward algebraic analysis in which we solve for the $\lambda_*$ value corresponding to $Q[\kappa] = 0$ for the give buckling mode curvature profile ansatz. Combining the curvature profile ansatz for a specific buckling mode with Eq. 3.56, we solve for that mode's critical buckling length above which the chain is no longer stable against buckling.

The ansatz for $\kappa(\zeta)$ must satisfy the boundary conditions in Eq. 3.54. To ensure the ansatz represents the actual curvature in buckled chains as closely as possible, we use simulations from the rigid model to inform our choice. Consider a simulation for $\Psi = 100$ and $a = 1$. Fig. 3.10A shows this simulation clearly exhibits the buckling instability marked by a smooth curvature profile through out the chain. Use the Menger curvature method, we compute the curvature at each linkage throughout the chain. We plot these curvature values against the parameterized location $\zeta$ in Fig. 3.10B. The curvature nearly satisfies the appropriate boundary conditions (though not exactly since the first and last linkages do not correspond to the chain tips at $\zeta = -1, 1$) and we observe the curvature vacillates between positive and negative extrema, with the largest curvature values occurring in the center of the chain at $\zeta = 0$. In the example shown in Fig 3.10, we observe x peaks in the curvature profile, and it is easy to imagine a curvature profile that has a similar form with either fewer or more peaks. The simplest example would be a curvature profile that is zero at $\zeta = -1$, increases as we approach the center of the chain, then decreases to zero at $\zeta = 1$ in a symmetric manner. This represents the first buckling mode ($n = 1$). Based these observations we devise the buckling curvature ansatz presented in Eq. 3.10. In our buckling mode ansatz the $n^{\text{th}}$ buckling mode curvature will have $n$ extreme points and since the simulation in Fig 3.10 shows five clear extremes in the curvature, we expect this chain is exhibiting the fifth buckling mode. We perform a least-square fit of the fifth buckling mode as shown in Fig 3.10 to the simulation data and find simulation and the mathematical ansatz are of similar form.

With the ansatz presented in Eq. 3.10, we can numerically solve for the $\lambda_*$ associated with each buckling mode. We plug the form of the $n^{\text{th}}$ buckling mode into Eq. 3.56, set it equal to zero and solve for $\lambda_*$. From the critical $\lambda_*$ values associated with each buckling mode, we can compute the critical buckling length associated with the emergence of each mode using Eq. 3.60.

### C Critical breaking stress coefficients

Eq. 3.13 provides the critical kinking length as a function of $\Psi$ and $a$. This critical length depends on two coefficients $\beta$ and $\gamma$ that vary with $a$. The coefficients are found during the perturbation analysis of the rigid rod model detailed in Appendix 2.B where they are referred to as $m$ and $z$. The coefficients have been renamed $\beta$ and $\gamma$ here and in Chapter 3 in order to avoid confusion with the bending modulus $m$ of the continuum mechanics framework. $\beta$ and $\gamma$ are found via linear regression of the critical breaking stress as a function of $\Psi$, but the process must be repeated for each desired value of $a$. The derivation in Appendix 2.A is for $a = 2$ but the process is easily repeated for any value of $a$. Since our simulations and analysis in Chapter 3 consider $a = 1, 2, 5, 10, 20, 50,$ and 100, we derive the coefficients $\gamma$ and $\beta$ for those specific values of $a$. The coefficients are presented in Table 3.2. Using these values, we obtain the non-dimensional critical breaking stress as a function of $\Psi$ used in Eqs. 3.13 and 3.16 for each of the desired values of $a$.

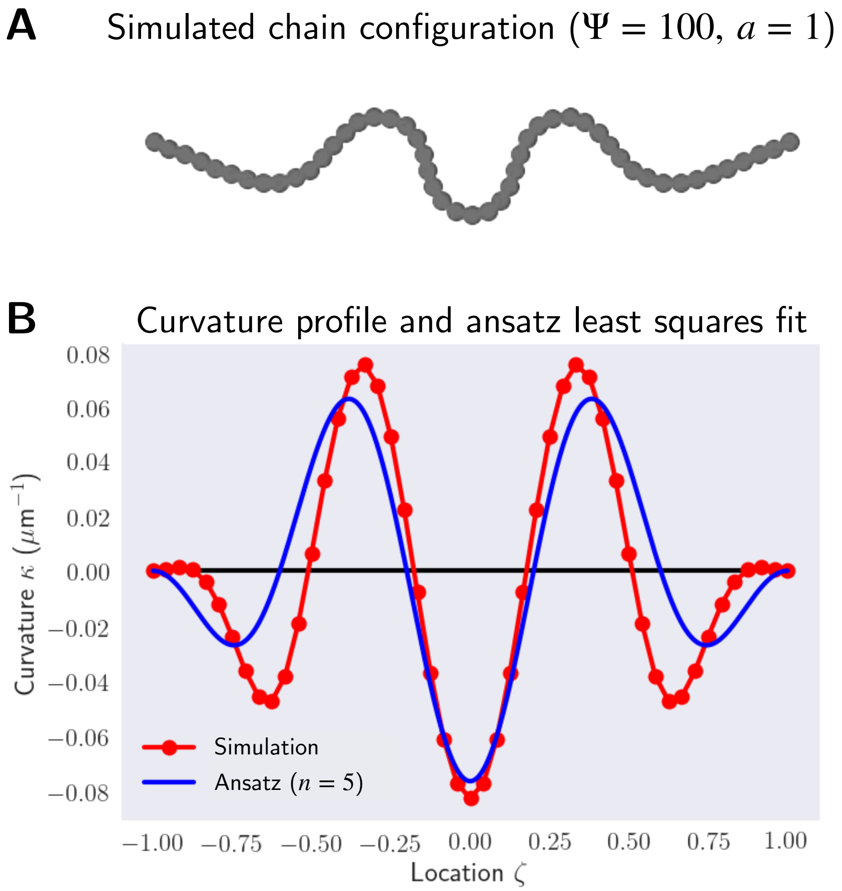


**Figure 3.10: Buckling curvature ansatz is based on simulation observations.** Simulation snapshot of a cell chain using the rigid rod model for $\Psi = 100$ and $a = 1$. This snapshot is from $t = 75$ mins from the the simulations shown in Fig 3.3. (A) The chain configuration, clearly exhibiting large curvatures as a result of buckling emergence. The points correspond to the linkages between cells. (B) The curvature profile throughout the chain as a function of the parameterized position $\zeta$ (red). The blue curve shows a least squares fit of the 5th buckling mode curvature as defined by Eq. 3.10.
